# RAPID RESISTANCE TO BET INHIBITORS IS MEDIATED BY FGFR1 IN GLIOBLASTOMA

**DOI:** 10.1101/2023.12.19.572401

**Authors:** Anna M. Jermakowicz, Alison M. Kurimchak, Katherine J. Johnson, Florence Bourgain-Guglielmetti, Simon Kaeppeli, Maurizio Affer, Hari Pradhyumnan, Robert Suter, Winston Walters, Maria Cepero, James Duncan, Nagi G. Ayad

**Author notes:** Corresponding author: Nagi G. Ayad.

## Abstract

Bromodomain and extra-terminal domain (BET) proteins are therapeutic targets in several cancers including the most common malignant adult brain tumor glioblastoma (GBM). Multiple small molecule inhibitors of BET proteins have been utilized in preclinical and clinical studies. Unfortunately, BET inhibitors have not shown efficacy in clinical trials enrolling GBM patients. One possible reason for this may stem from resistance mechanisms that arise after prolonged treatment within a clinical setting. However, the mechanisms and timeframe of resistance to BET inhibitors in GBM is not known. To identify the temporal order of resistance mechanisms in GBM we performed quantitative proteomics using multiplex-inhibitor bead mass spectrometry and demonstrated that intrinsic resistance to BET inhibitors in GBM treatment occurs rapidly within hours and involves the fibroblast growth factor receptor 1 (FGFR1) protein. Additionally, small molecule inhibition of BET proteins and FGFR1 simultaneously induces synergy in reducing GBM tumor growth *in vitro* and *in vivo*. Further, FGFR1 knockdown synergizes with BET inhibitor mediated reduction of GBM cell proliferation. Collectively, our studies suggest that co-targeting BET and FGFR1 may dampen resistance mechanisms to yield a clinical response in GBM.

## INTRODUCTION

Glioblastoma (GBM) accounts for 50 percent of patients with gliomas, making it the most common form of brain cancer ^1^. GBM currently has a median progression free survival of 6.9 months and median overall survival of 14.6 months ^2^. The standard of care for primary treatment, established in 2005, is surgical resection followed by radiation and temozolomide (TMZ) chemotherapy. However, GBM is fatal due to the diffuse infiltration of GBM cells into the surrounding brain tissue and the nearly universal resistance to both TMZ and radiation ^3,4^. GBM rapidly gains resistance to TMZ due to either an inherent or acquired overexpression of the O6-methylguanine DNA methyltransferase (MGMT) DNA repair, overexpression of epidermal growth factor receptor (EGFR), or restoration of p53 activity by Mdm2 inhibition ^5,6^. Although preliminary studies suggest that combination therapy with targeted agents may increase tumor radiosensitivity, ongoing clinical trials have shown minimal survival benefit ^7–9^. Additionally, due to toxicity limitations of radiotherapy, doses are insufficient to irradiate all tumor cells, which may promote tumor resistance and growth ^4,10^.

Furthermore, targeted therapies are largely ineffective in part due to intrinsic resistance pathways such as kinome reprogramming, which involves the dysregulation of kinase activity to trigger alternative survival pathways. Kinome reprogramming is thought to underlie the resistance of cancers to BET inhibitors, presenting a roadblock in the treatment of GBM. The BET family protein bromodomain-containing protein 4 (BRD4) is a therapeutic target in brain cancers, including GBM ^11–18^. BRD4 binds to acetylated lysine residues on histones and recruits and activates positive transcription elongation factor b (P-TEFb) complex to chromatin. P-TEFb then phosphorylates and activates RNA polymerase II (RNA Pol II) to initiate gene transcription. Acutely, inhibition of BRD4 blocks transcription of downstream oncogenes, including c-MYC, resulting in tumor reduction and apoptosis, however resistance is rapidly acquired ^19–22^. For example, in acute myeloid leukemia (AML), intrinsic resistance to BET inhibition results in activation of a c-MYC enhancer that compensates for the loss of BRD4 by utilizing WNT signaling to drive oncogenesis ^23^. Furthermore, several studies have examined the role of receptor tyrosine kinases (RTKs) in resistance to the BET inhibitor JQ1 ^24–26^. Recent studies have shown that kinome reprogramming may underlie this resistance by activating pro-survival RTKs, leading to compensatory pathways ^20,21^. This intrinsic drug resistance is important to consider in designing combination therapies ^27^. Therefore, combining BET inhibition and kinase inhibition is a potential avenue for eliciting the therapeutic effects of RTK inhibition while mitigating resistance mechanisms.

Fibroblast growth factor receptors (FGFRs) are comprised of a family of four receptor tyrosine kinases (RTKs) with a transmembrane domain, three extracellular immunoglobulin-like domains, and a split tyrosine kinase domain. The tyrosine kinase activity of FGFRs is activated by dimerization and autophosphorylation upon binding of the immunoglobulin-like domain to fibroblast growth receptor ligands and heparin sulfate proteoglycans ^28,29^. Activated FGFR leads to cellular proliferation, migration, angiogenesis, and reduced apoptosis via upregulation of RAS/MAPK, PI3K/AKT, and JAK/STAT pathways ^30,31^.

Here we report that resistance to BET inhibitors (BETi) is mediated by FGFR1, which is upregulated at the protein level within hours of BET protein inhibition. We profiled the kinome of GBM cells treated with the BET inhibitor JQ1 and we found a rapid upregulation of FGFR1 activity. Importantly, inhibition or knockdown of FGFR1 (FGFR1i) synergizes with the BET inhibitor JQ1, suggesting that FGFR1 signaling is an important resistance mechanism to BET inhibitors. This was also shown *in vivo* where BETi-FGFR1i combinations reduced tumor growth in mice relative to monotherapy. Collectively, our studies suggest that the rapid upregulation of FGFR1 is an important resistance mechanism to BET inhibitors in GBM.

## RESULTS

### Kinome profiling shows upregulation of key RTKs in response to BET inhibition

To explore the adaptive response of the kinome to BET protein inhibition after JQ1 treatment in GBM we applied quantitative MIB/MS kinome profiling technology. The results demonstrate that JQ1 induces kinome reprogramming as shown by PCA analysis and hierarchical clustering (Fig. 1a-b). Out of the 265 kinases detected using the MIB/MS profiling technology, 14 kinases (5%) were found to be significantly inhibited and 38 kinases (14%) were found to be significantly activated (Supplementary Table 1). Increased MIB-binding of RTKs EGFR, FGFR1, EPHA7, EPHB3 was observed, as well as downstream MEK-ERK-RSK1 signaling (MAP2K1, MAPK3 and RPS6KA1) and JNK signaling (MAP2K3, and MAPK8) in GBM22 cells following JQ1 treatment. Inhibition of cell cycle proteins CDK6, AURKA, and AURKB was shown by MIB/MS consistent with the observed cell cycle arrest (Fig. 1c).

**Figure 1.**
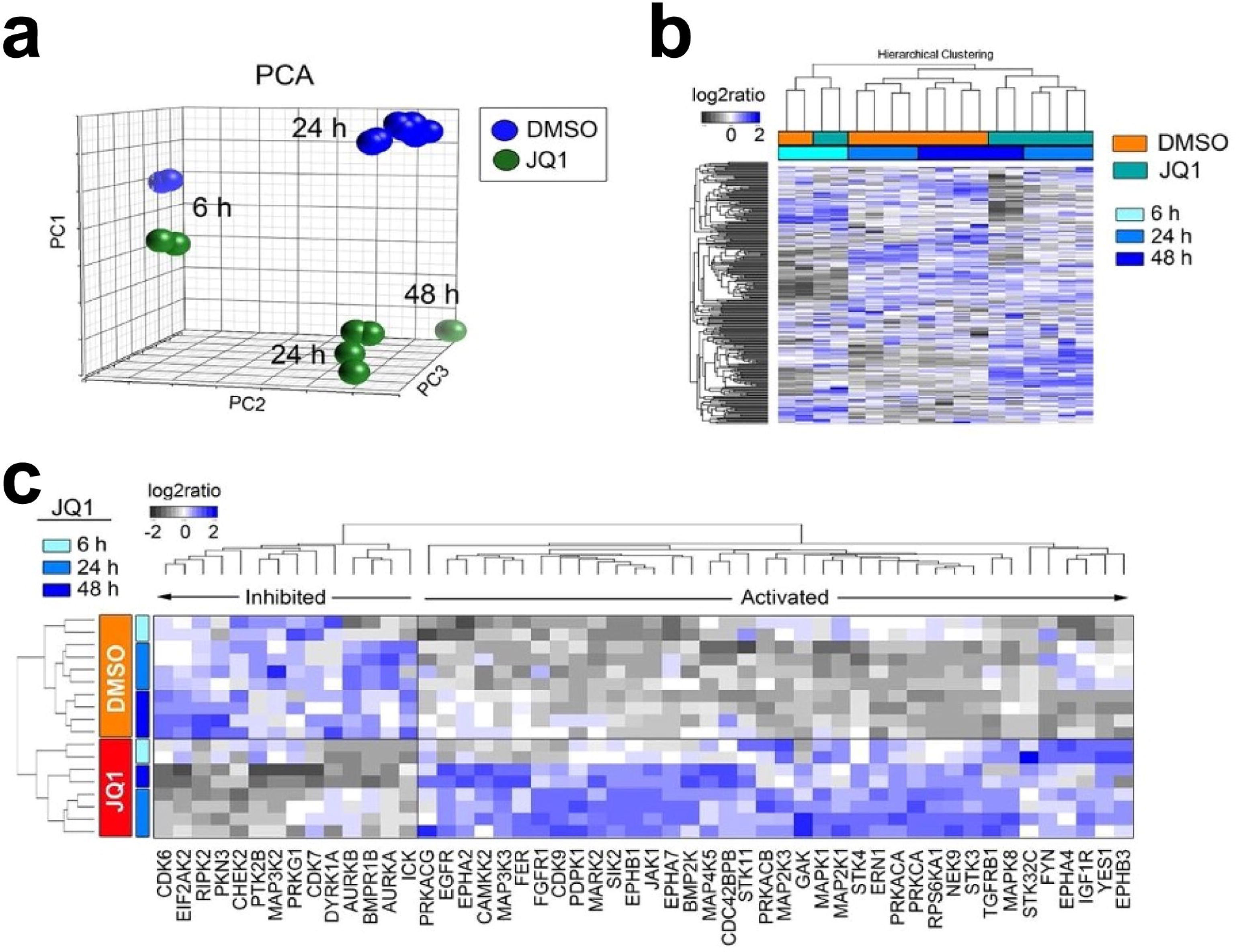
Dynamic adaptive reprogramming of the kinome to BET bromodomain inhibition in PDX-derived GBM cells. **a-b.** PCA analysis and unbiased hierarchical clustering of MIB/MS-defined GBM kinome in response to 6, 24 or 48 hours JQ1 therapy. **c.** Distinct statistical (ANOVA BH P≤0.05) log2 signature of kinases activated or repressed by JQ1 after 6, 24 or 48 hours compared to DMSO controls.

Protein network analysis of activated kinases revealed a tight cluster of MAPK activity and Ephrin receptor activity (Supplementary Fig. 1). KEGG pathway analysis also revealed an increase in autophagy, gonadotropin-releasing hormone (GnRH) signaling, and Ras signaling. FGFR1 has been previously shown to be essential for GnRH neurons, which promote cell proliferation and inhibit apoptosis in cancer cells ^32^. Interestingly, among the inhibited kinases we found a decrease in Aurora kinase A/B activity and pathway analysis revealed a decrease in the G1/S transition, indicating a possible arrest in the G1 phase as we saw previously with the BET inhibitor UM-002 (Supplementary Fig. 1) ^33^.

Since FGFR1 was found to be rapidly and transiently activated following BET inhibition (Fig. 1c), we focused on further analyzing this kinase as a potential early mediator to BET inhibitor resistance. FGFR1 has been shown to be a dependent gene when deleted in glioblastoma cells according to the Dependency Map ^34^ and FGFR1 mRNA expression correlates with poor survival in glioblastoma (Supplementary Fig. 2). Additionally, it has been demonstrated that FGFR1 mediates resistance to BET inhibitors in other cancers ^20^. Therefore, we assayed the protein levels of total FGFR1 following BET inhibition (Fig. 2a-b, Supplementary Figs. 3-5), as well as the alteration in gene expression by RT-qPCR (Fig. 2c). We found that total FGFR1 was increased at the protein level, despite no increase in gene expression, and this effect was diminished when BET inhibition was combined with FGFR inhibition.

**Figure 2.**
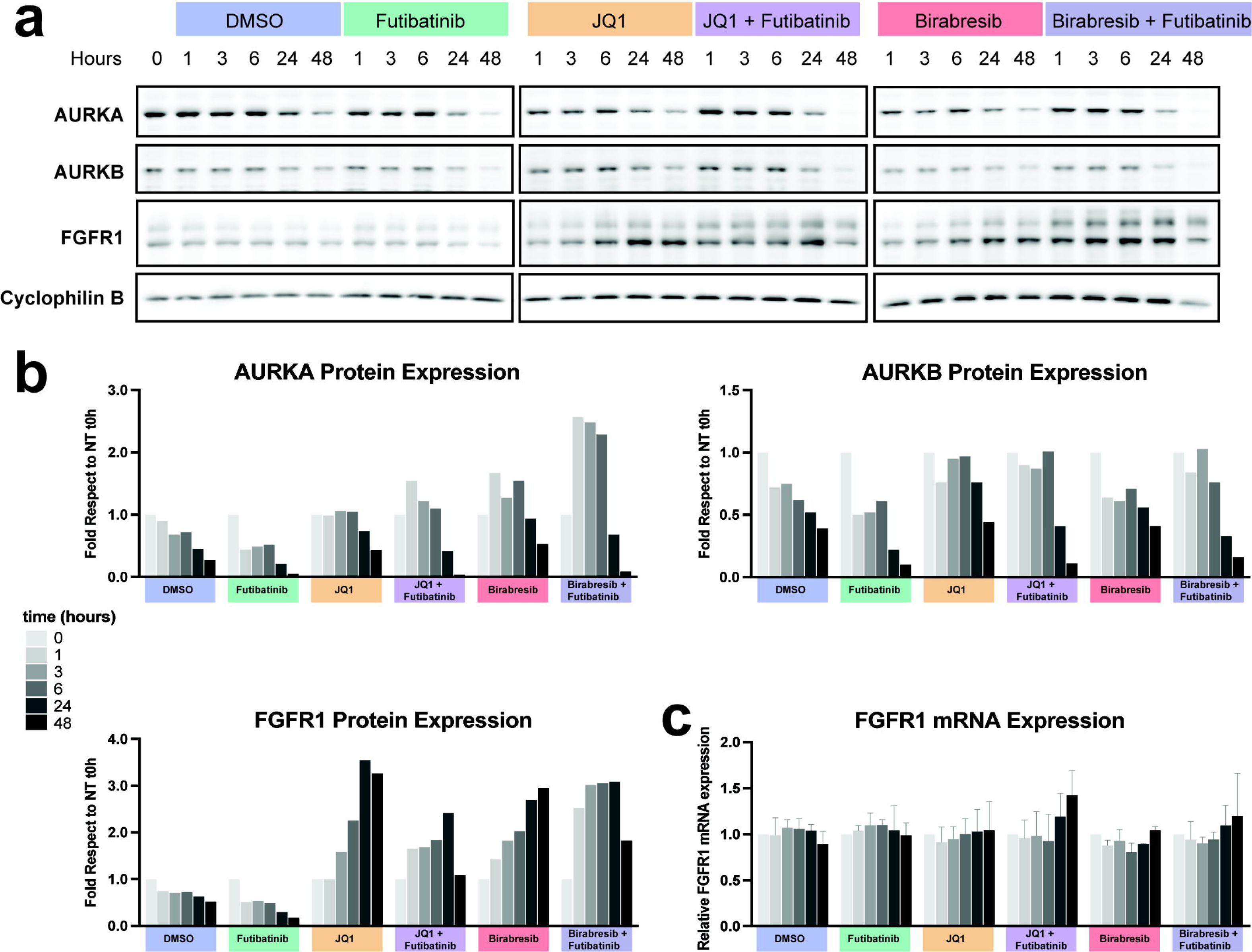
Expression of FGFR1 is increased in GBM22 cells treated with BET inhibitors. **a. FGFR1 and Aurora kinases expression.** GBM22 cells were treated with DMSO as a control or futibatinib, JQ1, JQ1+futibatinib, birabresib, birabresib+futibatinib, for 0, 1, 3, 6, 24, or 48 hours. Cell lysates were immunoblotted for FGFR1, Aurora Kinase A (AURKA), and Aurora Kinase B (AURKB). Equal loading was verified by immunoblotting for cyclophilin B from the respective gel. For cyclophilin B, the blot shown is from the same blot as FGFR1. The whole experiment was done in triplicate and one representative image is shown. **b. Quantification of protein expression from western blots in (a).** Quantification values for the target proteins were normalized to volumes of cyclophilin B loading control and plotted as fold difference from the time zero-hour point from the respective gel. Additional replicates can be found in Supplementary Figs. 3-4. **c. Quantification of FGFR1 mRNA expression.** FGFR1 mRNA level was assessed by RT-qPCR. For each time point, three PCR reactions were run and the average value was used for the calculation of relative expression using the ΔΔct method with GAPDH as a calibrator. Data are expressed as fold difference compared to the time zero hour. Graph shows the means and SD from three independent experiments.

### Synergy with combination therapy of BET inhibitors and an FGFR inhibitor *in vitro*

Based on the preliminary findings of FGFR1 protein activation in GBM following BET inhibition, we evaluated the synergistic potential of FGFR inhibitors with BET inhibitors. We tested futibatinib, a brain penetrant FGFR inhibitor, with the BET inhibitor JQ1 across a panel of newly diagnosed and recurrent GBM PDX lines. Compounds were combined in synergy matrices by overlapping the concentrations at each point. A highly synergistic response was seen using the Bliss independence model between different combinations of BET inhibitor and FGFR inhibitor (Fig. 3a-e, Fig. 4a-b). Importantly, each combination showed high synergy at low concentrations of each compound, which facilitates translation for *in vivo* studies since doses higher than 1 µM are typically not translatable to animal models without toxicity. The combination index of the Loewe additivity model was plotted in an isobologram, which revealed a highly synergistic effect as well. Additionally, FGFR1 knockdown was found to sensitize GBM cells to BET inhibition (Fig. 3f-g).

**Figure 3.**
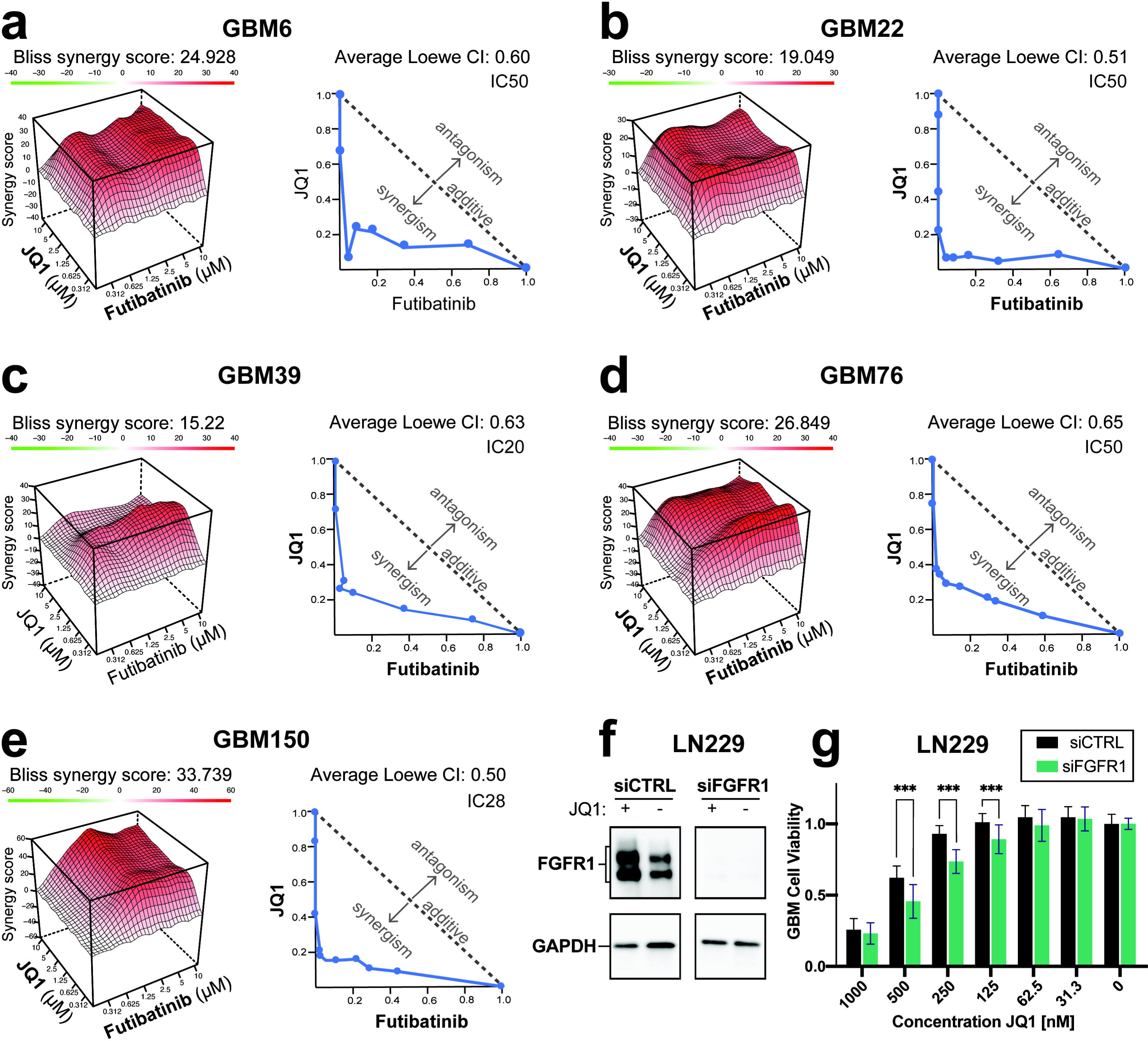
Combinations of BET inhibitor and FGFR inhibition in PDX GBM cells are synergistic in reducing cell proliferation. **a-e. Combinations of BET inhibition and FGFR inhibition in PDX GBM cells induces synergistic cell death.** GBM6, GBM22, GBM39, GBM76, and GBM150 cells were treated with a dose response matrix of futibatinib and JQ1. Cell death was measured as the amount of ATP present using CellTiter-Glo®. Synergy was assessed and visualized using the Bliss independence model synergy plot and Loewe additivity isobologram of the combination indices for each cell line at the indicated inhibition level. **f-g. Knockdown of FGFR1 sensitizes LN229 GBM cells to the BET inhibitor JQ1**. FGFR1 knockdown suppresses FGFR1 protein expression in LN229 cells (f). Following knockdown of FGFR1, treatment with JQ1 at decreasing concentrations was performed and the effect on cell viability was assessed (g). 16 replicates per group were tested, with significance determined using unpaired t-test and error bars calculated as standard deviation. Significance is represented by “*” where *, *P* < 0.05; **, *P* < 0.01; ***, *P* < 0.001.

**Figure 4.**
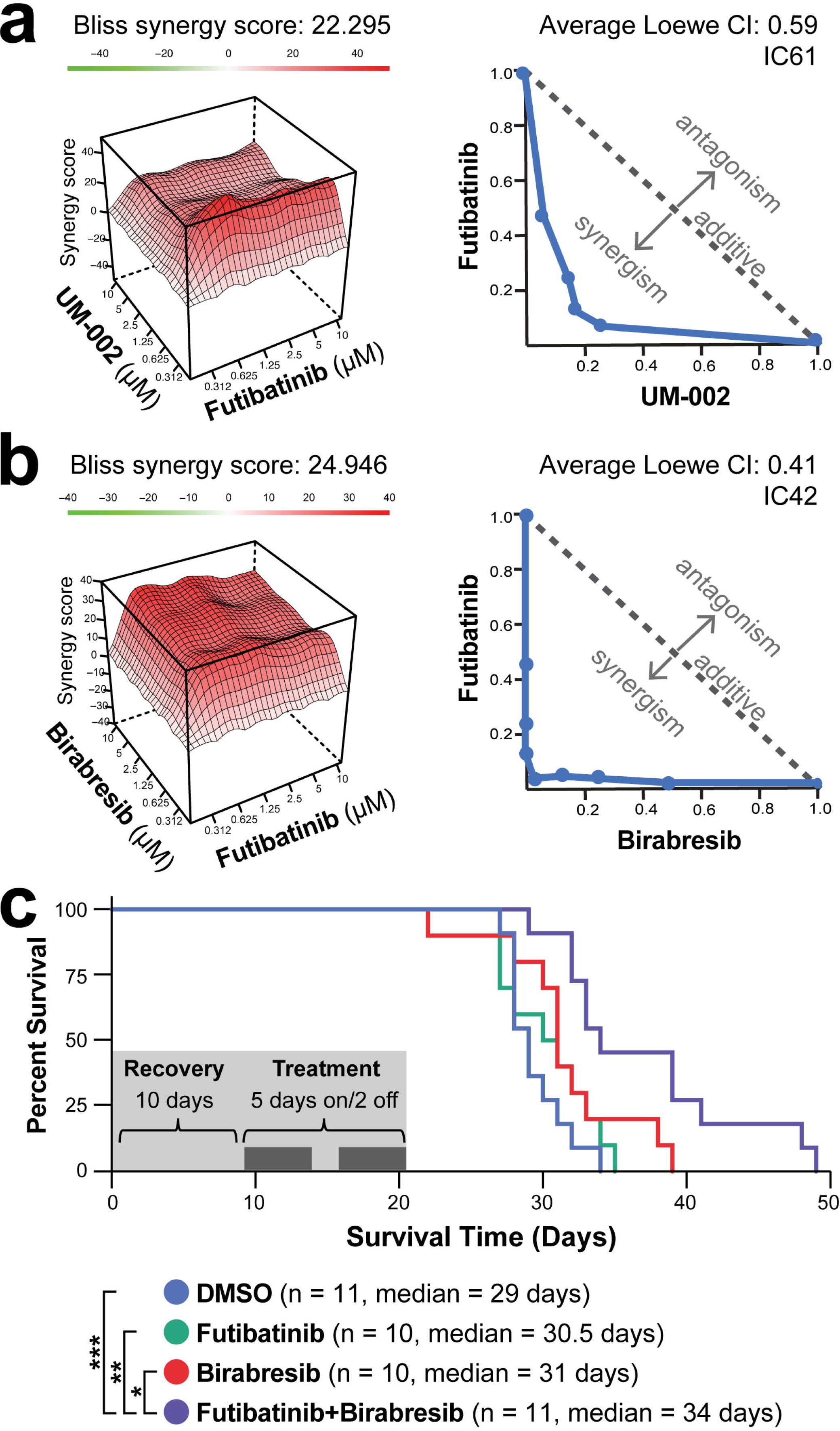
Combination of BET inhibitors and futibatinib induces synergy and improves survival in PDX GBM39. **a-c. In vitro combination screens show a synergistic increase in cell survival.** GBM39 cells were treated with a dose response matrix of the FGFR inhibitor futibatinib in combination with BET inhibitors (**a**) UM-002 or (**b**) birabresib. Cell death was measured as the amount of ATP present using CellTiter-Glo®. Bliss independence model synergy plot (top) and Loewe additivity isobologram (bottom) of the combination indices for each cell line at the indicated inhibition level. **c. Kaplan Meier curve of overall survival for mice bearing orthotopic GBM tumors treated with BET and/or FGFR inhibitors.** Following surgery (day 0), mice were given a 10 day recovery period before being assigned treatment groups. Mice were then treated with DMSO, birabresib, futibatinib, or birabresib + futibatinib for a total of 14 days (5 days on/2 days off). Animals were monitored for overall survival. Log-rank (Mantel-Cox) test was used to assess significance between the treatment groups. Significance is represented by “*” where *, P < 0.05; **, P < 0.01; ***, P < 0.005.

Since JQ1 is not a candidate for clinical trials and has shown limited efficacy in mouse models of GBM ^35,36^, we tested futibatinib in combination with additional BET inhibitors, namely UM-002 (Reaction Biology, Malvern, PA, USA) and birabresib (Selleck Chemicals, Houston, TX, USA) ^37^. UM-002 is a recently developed brain penetrant BET inhibitor that has been previously shown to reduce cell proliferation and invasion in GBM^33^. Birabresib is a BET inhibitor with a favorable safety profile that has been used in clinical trials for several solid tumors, including glioblastoma ^38^. BMS-986158 is also in clinical trials and has been shown to be brain penetrant ^39^. We performed synergy screens in GBM39 PDX cells and observed a strong degree of synergy across all cell lines for the combination of futibatinib and each BET inhibitor. Synergy was again assessed using the Bliss independence model and the Loewe combination index. Results were visualized using a Bliss synergy plot or a Loewe combination index isobologram (Fig. 4).

### Birabresib and futibatinib combination therapy shows a survival benefit *in vivo*

Since birabresib has entered clinical trials and has an established safety profile with favorable brain penetrance (Supplementary Fig. 6), we selected this inhibitor to use in combination with futibatinib *in vivo*. Mice received orthotopic intracranial implants with PDX GBM39 cells and were dosed with birabresib, futibatinib, birabresib + futibatinib, or DMSO for two weeks. We visualized results on a Kaplan Meier curve and assessed significance using a log-rank (Mantel-Cox) test (Fig. 4c). The combination treatment was significantly more effective than either drug alone, and was the only treatment group to reach a statistically significant increase in survival over the DMSO control. This supports our hypothesis that combination therapy may provide a more effective treatment for GBM than monotherapies.

At the time of euthanasia, mice were perfused, and tumors were isolated for RNA sequencing. Differentially expressed genes were determined using NOISeqBIO and the top ten upregulated and downregulated enriched biological processes for each treatment group were plotted on a heatmap (Supplementary Fig. 7) ^40,41^. The top genes, as determined by fold enrichment, were input into STRING for further pathway analysis and network visualization (Supplementary Fig. 7) ^42,43^.

## DISCUSSION

We have sought to identify the pathways mediating resistance to bromodomain inhibitors in glioblastoma by temporally assessing the activity of the kinome using multiplex-inhibitor bead mass spectrometry. We found that the activity of multiple kinases are downregulated while others are upregulated after treatment of cells with the BET inhibitor JQ1. One of the main upregulated kinases is FGFR1, which increases expression and activation at the protein level, but not at the mRNA level, after BET inhibitor treatment. This protein upregulation occurs very rapidly, as early as 6 hours after BET inhibitor treatment, suggesting it is an early event in mediating resistance to BET inhibitors. FGFR1 inhibition or knockdown synergizes with BET inhibition in reducing proliferation of GBM cells *in vitro*. Further, the FGFR1i-BETi combination of futibatinib and birabresib increases survival in an orthotopic mouse model of glioblastoma. Collectively, our studies suggest that FGFR1 significantly contributes to the emergence of resistance to BET inhibitors in GBM.

Although our studies are supported by findings in other cancers where FGFR1 upregulation has been observed after BET inhibition, such as uveal melanoma, breast cancer, or ovarian cancer cells ^44,45^, we are the first to demonstrate that this is an early event in the intrinsic resistance mechanism. Prior studies have shown the role that RTKs play in BET inhibitor resistance ^25,26^. The increase in FGFR1 occurs prior to cell cycle changes of treated cells or any proliferation changes as evidenced by reduced AURKA or AURKB levels (Figs. 1-2, Supplementary Figs. 3-5). This is distinct from previous studies that have shown the synergistic effect of BET inhibition with AURKA/B inhibition ^46,47^, suggesting that the efficacy of BET inhibitors is immediately countered by resistance mechanisms that include FGFR1 protein upregulation. However, the exact means through which FGFR1 protein is increased still needs to be elucidated and may require analysis of FGFR1 downstream pathways via phospho-proteomic analysis. We demonstrated that FGFR1 mRNA levels do not change and therefore posttranscriptional mechanisms are likely involved to upregulate FGFR1 protein after BET inhibition. Future studies should be performed to determine the roles of post transcriptional events in the intrinsic resistance mechanisms and upregulation of FGFR1 activation.

For acquired resistance, enriched biological processes following *in vivo* treatment with this combination showed that there was a decreased enrichment for regulation of long-term neuronal plasticity, which may indicate that the enrichment of OPC-like cells is diminished with the combination treatment. Several of these pathways involve exosome activity, which have been implicated in signaling between the tumor and microenvironment ^48^. However, in both the birabresib and combination treatment groups we see an upregulation in angiogenesis-related cell migration, suggesting that a resistance mechanism could be related to increased endothelial growth factors. Ultimately, the upregulated biological processes point to cellular migration as a possible mechanism of tumor growth following treatment with our combination therapy (Supplementary Fig. 7).

Future studies are also essential for determining the BETi-FGFR1i combinations to be used in GBM clinical trials. We used two clinical compounds birabresib and futibatinib that showed a statistically significant improvement of survival of mice implanted orthotopically with GBM39 cells. However, the *in vivo* results were not as robust as the *in vitro* synergy we observed. There are multiple possible reasons for this finding. For one, we observed that whereas futibatinib was brain penetrant (14% brain:plasma ratio) in mice, birabresib was not as strongly penetrant (1% brain penetrant in non-tumor bearing, 6% penetrant in tumor-bearing mice), and therefore it is possible that not enough BET inhibitor reached the tumor cells within the brain (Supplementary Fig. 6). We have developed the more brain penetrant BET inhibitor UM-002 (11% brain penetrant) ^33^, which does synergize with futibatinib *in vitro* (Fig. 4a). However, combination treatment with UM002 + futibatinib was not well tolerated. Several promising BET inhibitors have recently entered clinical trials. Trotabresib is in a phase 1 clinical trial for high grade gliomas and has been shown to cross the blood brain barrier effectively ^49^. Our preliminary studies also suggest that this inhibitor synergizes with futibatinib (Supplementary Fig. 8) and therefore new BET inhibitors should be considered as a potential therapeutic avenue for GBM combination treatment with futibatinib.

Another possible reason for reduced efficacy of the BETi-FGFR1i combination *in vivo* is that resistance mechanisms remain even when using two inhibitors. This suggests that the initial intrinsic resistance observed to JQ1 in the kinome screen may in fact be overcome by FGFR1 inhibition, however acquired resistance is eventually obtained to the combination treatment. Our RNA sequencing studies from treated mice support this since pathways involved in migration and invasion are upregulated with the birabresib/futibatinib combination (Supplementary Fig. 7). Future studies are required to determine whether co-targeting of these pathways significantly improves GBM patient survival in combination with birabresib/futibatinib. Importantly, we have demonstrated that when assessing these future combinations within a clinical setting it is essential to consider the involvement of FGFR1 resistance mechanisms at the early stages of GBM treatment.

## METHODS

### Cell culture conditions

Patient-derived xenograft (PDX) cells GBM6, GBM22, GBM39, GBM76, and GBM150 were obtained from the Brain Tumor PDX national resource at the Mayo Clinic^50^. Cells were cultured in complete media consisting of Dulbecco’s Modified Eagle’s medium (DMEM):F12 with 10% fetal bovine serum and 1% penicillin and streptomycin at 37°C in 5% CO2 and saturated moisture. Cells were maintained for a maximum of 30 days before being discarded ^51^. The diagnostic status and molecular features of the PDX cells used are outlined in Supplementary Table 2.

LN229 cells (obtained from ATCC) were cultured in DMEM:F12 medium (Gibco, Life Technologies, Carlsbad, CA, USA) with 5% fetal bovine serum (Gibco, Life Technologies, Carlsbad, CA, USA) and 1% Penicillin-Streptomycin (Gibco, Life Technologies, Carlsbad, CA, USA) at 37°C in 5% CO2 and saturated moisture. Cells were regularly tested to ensure absence of mycoplasma and underwent IMPACT testing before being used in animal experiments.

### Kinome profiling of PDX GBM22 cells after BET inhibition using multiplexed kinase inhibitor beads and quantitative mass spectrometry (MIB-MS)

We profiled the kinome in GBM PDX22 cells before and after incubation with JQ1 or DMSO for 6, 24, or 48 hours. PDX GBM22 cells were cultured in dishes and grown to 80% confluency. JQ1 (Selleck Chemicals, Houston, TX, USA) was added to media for a final concentration of 500 nM and cells were harvested at 0 hours, 6 hours, 24 hours, and 48 hours. Cells were extracted following an established protocol ^20,52^. Cells were lysed on ice in buffer containing 50 mM HEPES (pH 7.5), 0.5% Triton X-100, 150 mM NaCl, 1 mM EDTA, 1 mM EGTA, 10 mM sodium fluoride, 2.5 mM sodium orthovanadate, 1X protease inhibitor cocktail (Roche, Basel, Switzerland), and 1% each of phosphatase inhibitor cocktails 2 and 3 (Sigma, Saint Louis, MO, USA). Particulate was removed by centrifugation of lysates at 13,000 rpm for 10 minutes at 4°C and filtration through 0.45 µm syringe filters and lysates were stored at -80°C until preparation for MIB-MS. Kinase extracts were analyzed using SILAC mass spectrometry following a previously established protocol ^20,52^.

An equal amount of s-SILAC reference ([13C6, 15N4 ] arginine (Arg 10 ) and [ 13C6 15N2 ] lysine (Lys 8 )) (5 mg) lysate was added to non-labeled (5 mg) sample and analyzed on MIB-beads. Endogenous kinases were isolated by flowing lysates over kinase inhibitor-conjugated Sepharose beads (purvalanol B, VI16832, PP58 and CTx-0294885 beads) in 10 mL gravity-flow columns. After 2×10 mL column washes in high-salt buffer and 1×10 mL wash in low-salt buffer (containing 50 mM HEPES (pH 7.5), 0.5% Triton X-100, 1 mM EDTA, 1 mM EGTA, and 10 mM sodium fluoride, and 1 M NaCl or 150 mM NaCl, respectively), retained kinases were eluted from the column by boiling in 2×500 µL of 0.5% SDS, 0.1 M TrisHCl (pH 6.8), and 1% 2-mercaptoethanol. Eluted peptides were reduced by incubation with 5 mM DTT at 65°C for 25 minutes, alkylated with 20 mM iodoacetamide at room temperature for 30 minutes in the dark, and alkylation was quenched with DTT for 10 minutes. Samples were concentrated to approximately 100 µL with Millipore 10kD cutoff spin concentrators. Detergent was removed by methanol/chloroform extraction, and the protein pellet was resuspended in 50 mM ammonium bicarbonate and digested with sequencing-grade modified trypsin (Promega, Madison, WI, USA) overnight at 37°C. Peptides were cleaned with PepClean C18 spin columns (Thermo Fisher Scientific, Waltham, MA, USA), dried in a speed-vac, resuspended in 50 μL 0.1% formic acid, and extracted with ethyl acetate (10:1 ethyl acetate:H2O). Briefly, 1 mL ethyl acetate was added to each sample, vortexed and centrifuged at max speed for 5 minutes, then removed. This process is repeated 4 more times. After removal of ethyl acetate following the 5th centrifugation, samples were placed at 60°C for 10 minutes to evaporate residual ethyl acetate. The peptides were dried in a speed vac, and subsequent LC-/MS/MS analysis was performed. Proteolytic peptides were resuspended in 0.1% formic acid and separated with a Thermo RSLC Ultimate 3000 on a Thermo Easy-Spray C18 PepMap 75µm x 50cm C-18 2 µm column with a 240 min gradient of 4-25% acetonitrile with 0.1% formic acid at 300 nL/min at 50°C. Eluted peptides were analyzed by a Thermo Q Exactive plus mass spectrometer utilizing a top 15 methodology in which the 15 most intense peptide precursor ions were subjected to fragmentation. The AGC for MS1 was set to 3×10^6^ with a max injection time of 120 ms, the AGC for MS2 ions was set to 1×10^5^ with a max injection time of 150 ms, and the dynamic exclusion was set to 90 s. Raw data analysis of SILAC experiments was performed using MaxQuant software version 1.5.3.30 and searched against the Swiss-Prot human protein database (downloaded on September 25, 2015). The search was set up for full tryptic peptides with a maximum of two missed cleavage sites. All settings were default and searched using acetylation of protein N-terminus and oxidized methionine as variable modifications. Carbamidomethylation of cysteine was set as fixed modification. SILAC quantification was performed by choosing multiplicity as 2 in group-specific parameters and Arg10 and Lys8 as heavy labels. Match between runs was employed and the significance threshold of the ion score was calculated based on a false discovery rate of < 1%. MaxQuant normalized ratios were analyzed as follows: for a total of p unique kinases, we computed the pooled protein ratio and p-value across the replicates. For each replicate, we identified kinases that exhibit statistically significant changes in expression based on step-up adjusted p-values at FDR of 0.05 to account for multiple comparisons. Principal component analysis (PCA) of kinome profiling MIBs-values was performed using Partek Genomics Suite (Partek Inc.). Visualization of MIBs-kinome signatures using PCA plots or Log 2 ratio heat maps were generated using Partek Genomics Suite.

### Western blot analysis

Cells were cultured as described previously with the BET inhibitors JQ1 or birabresib in combination with the FGFR inhibitor futibatinib at doses sufficient to induce up to 50% cell death ^53,54^. Compounds used to treat cells were dissolved in 100% dimethyl sulfoxide (DMSO, Sigma, Saint Louis, MO, USA). Cells were treated with DMSO as a control, 500 nM JQ1, 1000 nM birabresib (MedChemExpress, Monmouth Junction, NJ, USA), and/or 250 nM futibatinib (MedChemExpress, Monmouth Junction, NJ, USA). Cells were harvested at 1, 3, 6, 24 and 48 hours after the treatments. Cells were rinsed with PBS and then lysed with the Western Lysis Buffer (1:10, PhosphoSolutions, Aurora, CO, USA) supplemented with protease/phosphatase inhibitor cocktail (1:100, Cell Signaling Technologies, Danvers, MA, USA). The lysates were kept on ice for 10 min before being sonicated 3 times for 5 seconds. Lysates were then centrifuged for 20 min at 17,200 RCF at 4°C to remove any cellular debris. The supernatant was collected, and lysate protein concentrations were determined by a BCA protein assay (Thermo Scientific Pierce, Waltham, MA, USA). Equal amounts of protein (15 to 20 μg for each lane) were separated by SDS-PAGE using Novex WedgeWell 4-20% Tris-Glycine gels (Fisher Scientific, Waltham, MA, USA) and transferred to nitrocellulose membranes (Protran 0.1 μm, Amersham Biosciences, Amersham, UK). Immunoblots were blocked with 5% BSA in TBS-Tween 20 (0.05%, v/v) for 1 hour at room temperature. Membranes were then incubated with primary antibodies diluted in 2.5% BSA in TBS-Tween 20 (0.05%, v/v), overnight at 4°C. The antibodies used are outlined in Supplementary Table 3.

Following several washes with TBS-Tween 20 (0.05%, v/v), each membrane was incubated with a secondary antibody conjugated with horseradish peroxidase (Cell Signaling Technologies, Danvers, MA, USA), diluted in 2.5% BSA in TBS-Tween 20 (0.05%, v/v) for 90 minutes at room temperature. The blots were developed using an enhanced chemiluminescence western blotting detection system (SuperSignal West Dura Extended Duration Substrate, Thermo Scientific, Waltham, MA, USA). Images were acquired with a digital imager (Azure Biosystems c600, Dublin, CA, USA). Band density was quantified using Image Studio Software from LI-COR Biosciences (Lincoln, NE, USA). Quantification values for the target proteins were normalized to values of Cyclophilin B loading control and plotted as fold difference from the non-treated time zero-hour point. Graphical representation of the quantification data was created using GraphPad Prism 9.

### RT-qPCR

An RT-qPCR assay was used to determine the level of *FGFR1* expression after each single compound or combination treatment. Cells were cultured as described previously and treated with DMSO as a control, 500 nM JQ1 or 1000 nM birabresib, alone or in combination with 250 nM futibatinib for 0, 1, 3, 6, 24, or 48 hours. The cDNA was prepared using SuperScript III (ThermoFisher, Waltham, MA, USA) and random primers following the manufacturer’s instructions: 2 µg of RNA were retro-transcribed in 20 µL, volume was then increased to 40 µL and 4 µL were used for each PCR reaction.

PowerUp SYBR Green master mix (Applied Biosystems, Waltham, MA, USA) was used with the following conditions: 2 minutes at 50°C, 2 minutes at 95°C, 15 seconds at 95°C, and 1 minute at 60°C. All samples were run in triplicate on a Quant Studio 6 Flex machine (Applied Biosystems, Waltham, MA, USA) and analyzed using QuantStudio Design and Analysis Software (Applied Biosystems, Waltham, MA, USA). The relative expression of FGFR1 was calculated using the ΔΔCt method with GAPDH as a reference. Forward and reverse primers were designed using Primer 3 software (available online at: https://bioinfo.ut.ee/primer3-0.4.0/) (Supplementary Table 4).

### FGFR1 siRNA knockdown and JQ1 treatment

LN229 cells were seeded in opaque white 96-well plates at 6,000 cells per well and treated the following day. siRNA knockdown was performed either with siFGFR1 (ON-TARGETplus Human FGFR1 siRNA SMART Pool, Horizon Discovery, Cambridge, UK) or non-targeting siControl (ON-TARGETplus Non-targeting Control Pool, Horizon Discovery, Cambridge, UK) using Lipofectamine RNAiMAX transfection reagent (Invitrogen, Waltham, MA, USA). Briefly, the transfection complexes were generated according to the manufacturer’s guidelines and then added to 1.5 mL tubes containing antibiotics-free medium with 0.1% DMSO to a final siRNA concentration of 17 nM. Cells were incubated with or without 500 nM of JQ1 for 5 days prior to protein extraction. Cell lysates were prepared as described above. Nitrocellulose membranes were blocked for 1h at RT in 5% Blotting Grade Blocker Non-Fat Dry Milk (Biorad, Hercules, CA, USA) and TBST. All antibodies were diluted 1:1000 in 5% milk/TBST (FGFR1: D8E4; GAPDH HRP-conjugated: 14C10; Cell Signaling, Danvers, MA, USA). The membranes were incubated with antibodies at 4°C overnight. The FGFR1-probed membranes were washed 5x for 5min in TBST and then probed with anti-rabbit HRP-conjugated antibody (7074S, Cell Signaling, Danvers, MA, USA) for 1h at RT. All membranes were then washed five times in TBST and signal was acquired using the SuperSignal West Femto Maximum Sensitivity chemiluminescence substrate (Fisher Scientific, Hampton, NH, USA) as described above.

Finally, a 1:2 dilution series of JQ1 (MedChemExpress, Monmouth Junction, NJ, USA) ranging from 1000 nM to 1.95 nM final concentration was performed, including a 0.1% DMSO (Tocris Bioscience, Bristol, UK) and 10 μM Velcade (Selleck Chemicals, Houston, TX, USA) ^55^ control in four replicates each. The cells were incubated with the siRNA/JQ1 treatment for five days. A CellTiter-Glo® Luminescent Cell Viability Assay (Promega Corporation, Madison, WI, USA) was carried out according to the manufacturer’s instructions. The plates were read in a CLARIOstar plate reader (BMG Labtech, Cary, NC, USA) using the luminescence protocol. The results were evaluated in Microsoft Excel (Microsoft, Redmond, WA, USA) and GraphPad Prism (Version 9.5.1, GraphPad Software, Boston, MA, USA).

### *In vitro* synergy screens for BET inhibition and FGFR inhibition

PDX cells were plated in 25 µL of complete media in Nunc® 384-Well Tissue Culture Plates (Thermo Scientific, Waltham, MA, USA) at a concentration of 3000 cells per well. Cells were incubated overnight to establish adherent cultures, treated with 5 µL of compound (at 6 times the final concentration) dissolved in DMSO and Hank’s Balanced Salt Solution, and then incubated for 72 hours. Finally, ATP content was measured using the CellTiter-Glo® Luminescent Cell Viability Assay (Promega Corporation, Madison, WI, USA) or the Caspase Glo® 3/7 Assay (Promega Corporation, Madison, WI, USA) following the manufacturer’s protocol and plates were read on an EnVision Multilabel Plate Reader (Perkin Elmer, Waltham, MA, USA). Synergy screens consisted of a minimum of three replicates of 7×7 dose-response matrices, ranging from 10 - 0.3125 μM at 1:2 dilutions with seven replicates each of DMSO as a negative control and 10 μM Velcade as a positive control. Final DMSO concentration was maintained at 0.2% in all treatment conditions. Reduced cell proliferation was measured by normalizing the raw fluorescent values to the negative control (DMSO, 0% reduction) and the positive control (Velcade, 100% reduction) using the following formula:

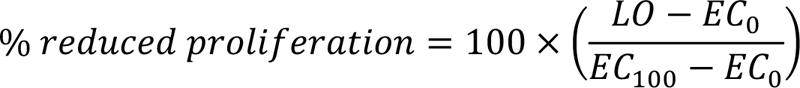

where LO is the raw luminescent output value, EC_0_ is the mean raw luminescent of the negative control, and EC_100_ is the mean raw luminescent output of the positive control. Each condition includes a minimum of three technical replicates, with a standard deviation less than 5.

To assess the synergy of the two drug treatments, two different graphs were produced: an isobologram of Loewe additivity model combination index and a 3D representation of the response surface of the combination matrix as quantified by the Bliss score.

The Loewe additivity model divides the amount of drug needed in combination therapy to achieve a given effect level by the amount of drug in monotherapy needed to achieve that same effect ^56^. If the addition of this metric for both drugs is less than 1, then the combination is said to be synergistic above the amount expected by adding the drugs together under the Loewe model. Combination index (CI) points are calculated using the following equation:

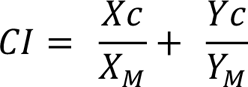

For the dose of drugs (X) and (Y) needed to achieve the specified effect level in monotherapy (M) or combination (C). A four-parameter dose-response curve was calculated in PRISM using the results from the 7×7 synergy matrices. The effect level (IC) for visualization on an isobologram was selected as the maximum effect level able to be regressed for both drugs in monotherapy, up to the IC_50_ level. Each point on the isobologram indicates the combination index for drug X and drug Y at each variable concentration ratio in the 7×7 matrix, where synergistic combinations will have a combined CI for drug X and drug Y below 1, as indicated by the dashed line.

The Bliss independence model is another widely utilized model that is based on the principle that drugs act independently and do not interfere with each other ^57^. It is calculated based on the formula below:

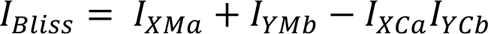

For drugs (X) and (Y) in monotherapy (M) or combination (C) at concentrations (a) and (b) for the inhibition level (I). if I_Observed_ > I_Bliss_ this indicates synergism, while I_Observed_ < I_Bliss_ indicates antagonism. Bliss independence scores were calculated and visualized using the synergyfinder package ^58^.

### *In vivo* orthotopic intracranial glioblastoma tumor implantation

Forty-four Nu/nu mice were implanted with 300,000 PDX GBM39 cells suspended in PBS using a previously published protocol ^51^. Mice were anesthetized with ketamine (100 mg/kg) and xylazine (10 mg/kg) and placed in a stereotactic frame with a mouse adaptor (Stoeling lab standard #51615). The site of incision was disinfected with Nolvasan and a 1 cm incision was made at the midline from the level of the eyes to the ears. After exposing the skull, bregma was visualized and a point 1 mm anterior and 2 mm lateral was marked. The skull was carefully drilled using a Dremel tool with a #8 bit. Using a 26 G Hamilton syringe (Hamilton, Reno, NV), 3 µL of cell suspension (100,000 cells/µl) was drawn up and the Hamilton syringe was mounted to a stereotaxic injector (Stoeling quintessential stereotaxic injector #53311). The needle was advanced to a depth of 3 mm, withdrawn to a depth of 2 mm, then cells were injected slowly over a period of 1 minute. After injection, the needle was held in place for an additional minute, then withdrawn. The scalp was sealed with surgical glue and sutured, treated with triple antibiotic, and injected with buprenorphine SR-Lab 1 mg/mL for pain management. Mice recovered for a period of 10 days before treatment.

### *In vivo* brain penetrance of inhibitors

Mice were treated with 50 mg/kg birabresib or 12.5 mg/kg futibatinib. Compounds were dissolved in DMSO and diluted to a final concentration of 10% DMSO, 40% PEG400, 50% PBS and injected intraperitoneally with a final volume of 10 µL/g body weight for three mice per group. 120 minutes after treatment, mice were anesthetized with 250 mg/kg avertin. Brains were isolated and separated into tumor and non-tumor hemispheres. Compound levels were determined by mass spectrometry using an AB Sciex 5500 mass spectrometer (Sciex, Toronto, CA) using multiple reaction monitoring by comparison against separate standard curves prepared in blank mouse brain homogenate.

### *In vivo* GBM tumor treatment and monitoring

Mice were treated with DMSO, 50 mg/kg/day birabresib, 12.5 mg/kg/day futibatinib, or 50 mg/kg/day birabresib + 12.5 mg/kg/day futibatinib. Compounds were dissolved in DMSO and diluted to a final concentration of 10% DMSO, 40% PEG400, 50% PBS. Compounds were injected intraperitoneally 5 days per week on alternating sides, with a final volume of 10 µL/g body weight. Drug treatment was administered for a duration of 2 weeks.

Mice that died early due to surgical complications were omitted from the study, leaving us with a final number of 10 mice per group for futibatinib or birabresib monotherapy, and 11 mice per group for DMSO or birabresib+futibatinib. Mice were monitored daily for signs of physical decline. At the time of death, tumors were collected for RNA sequencing.

### Isolation of tumors for RNA sequencing

Mice were anesthetized with tribromoethanol (avertin, 250 mg/kg Sigma-Aldrich) intraperitoneally prior to transcardial perfusion. Once the mouse showed signs of proper depth of anesthesia (lack of reflex response to hind paw withdrawal or ocular stimulation), the chest cavity was opened and the heart was exposed. A cut was made in the right atrium to prevent blood recirculation and a 28 G butterfly needle attached to a syringe was inserted into the left ventricle to perfuse the circulatory system with PBS. After perfusion, mice were rapidly decapitated and tumors were isolated from the brain. Tumors were dissociated using a mortar and pestle, followed by a QIAshredder column (Qiagen, Valencia, CA, USA). RNA was extracted from the lysate per manufacturer instructions using a AllPrep DNA/RNA mini kit (Qiagen, Valencia, CA, USA).

### RNA sequencing and gene expression analysis

RNA was isolated from nine samples for RNA sequencing: 2 samples for groups birabresib, futibatinib, and birabresib+futibatinib, and 3 samples from the DMSO control group. Tumor samples were sequenced on the NovaSeq 6000 (Illumina, San Diego, CA, USA). FastQ files were aligned to the human genome and raw gene counts were determined. Trimmed Mean of M-values (TMM) normalization was performed using edgeR ^59–61^. TMM values were used as input into the NOISeqBIO R-package for differential gene analysis ^40,41^. Genes with low counts across all samples were excluded, samples were normalized to the DMSO control, and differentially expressed genes were detected. Significantly enriched pathways were identified using EnrichR (https://maayanlab.cloud/Enrichr/) ^62,63^, including biological processes, gene ontology, and KEGG pathway analysis.

## ACKNOWLEDGMENTS

We would like to thank all members of the Ayad and Duncan laboratories for helpful discussions. We acknowledge funding from the BellRinger from Lombardi Comprehensive Cancer Center.

## AUTHOR CONTRIBUTIONS

A.M.J., A.M.K., S.K., J.S.D., and N.G.A. were responsible for conception, design, and developing methodology. A.M.J., R.K.S., S.K., and N.G.A. wrote the manuscript. A.M.J., R.K.S., S.K., A.M.K., K.J.J., M.A., F.B-G., H.P., W.W., and M.C. were responsible for data collection and acquisition. K.J.J., A.M.K., and J.S.D. prepared Fig. 1. F.B-G., H.P., and M.A. prepared Fig. 2. All authors reviewed and edited the manuscript.

## DATA AVAILABILITY STATEMENT

The raw datasets generated and analyzed for RNA sequence analysis of PDX GBM tumors treated with BET and/or FGFR inhibitors *in vivo* have been deposited in NCBI GeoDataset as GSE245624. The mass spectrometry proteomics data have been deposited to the ProteomeXchange Consortium via the PRIDE ^64^ partner repository with the dataset identifier PXD043214.

## ADDITIONAL INFORMATION

The authors declare no competing interests.

