## Supplementary Files for "RAPID RESISTANCE TO BET INHIBITORS IS MEDIATED BY FGFR1 IN GLIOBLASTOMA"

**SUPPLEMENTARY INFORMATION**

**Supplementary Table 2. PDX GBM cell line diagnostic status and molecular features.**

| PDX Cell Line | GBM Diagnosis | Subtype | EGFR | MGMT | IDH |
|---|---|---|---|---|---|
| GBM6 | Newly diagnosed | Classical | VIII | Unmethylated | Wildtype |
| GBM22 | Newly diagnosed | Classical | Wildtype | Methylated | Wildtype |
| GBM39 | Newly diagnosed | Mesenchymal | VIII | Methylated | Wildtype |
| GBM76 | Recurrent | Classical | VIII | Methylated | Wildtype |
| GBM150 | Recurrent | NA | Wildtype | Unmethylated | Wildtype |

**Supplementary Table 3. Western blot antibodies.**

| Antibody | Brand | Catalog # | Dilution |
|---|---|---|---|
| FGF Receptor 1 (D8E4) | Cell Signaling | 9740 | 1:500 |
| Phospho-FGF Receptor 1 (Tyr653/654) | Cell Signaling | 52928 | 1:500 |
| Aurora B/AIM1 | Cell Signaling | 3094S | 1:500 |
| Aurora A (D3V7T) | Cell Signaling | 91590 | 1:500 |
| Cyclophilin B (D1V5J) | Cell Signaling | 43603 | 1:10,000 |

**Supplementary Table 4. RT-qPCR primers.**

| Name | Sequence | Target gene | Genomic Position | Length | TM |
|---|---|---|---|---|---|
| GAPDH for | AACAGCCTCAAGATCATCAGC | GAPDH for | chr12:6,537,197-6,537,216 | 21 | 63 |
| GAPDH rev | GGATGATGTTCTGGAGAGCC | GAPDH rev | chr12:6,537,661-6,537,681 | 20 | 62 |
| o2 | GTCTTTCTCTGTTGCGTCCG | FGFR1 for | chr8:38,417,397-38,417,416 | 20 | 59 |
| o3 | CCAACCGTGTGACCAAAGTG | FGFR1 rev | chr8:38,417,889-38,417,908 | 20 | 59 |

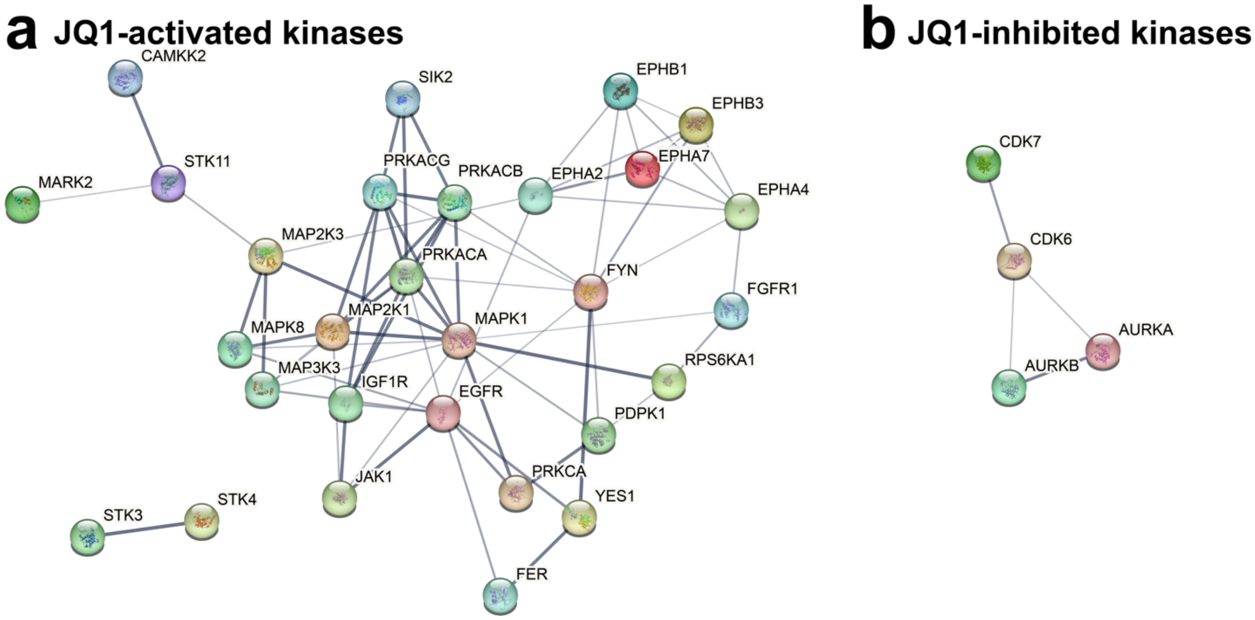


**Supplementary Figure 1. STRING protein networks reveal dysregulated signaling pathways in GBM cells after BET inhibition.** Differential MIB-binding of kinases was determined by an ANOVA, BH P≤0.05 and statistically significant kinases uploaded to the STRING database (https://string-db.org/ version 12.0) for further analysis. Kinase protein interactions with medium confidence interactions based on experiments, databases, co-expression, neighborhood, gene fusion, and co-occurrence were mapped using k-means clustering. Disconnected nodes in the networks were hidden. **a.** Protein network of statistically activated kinases. **b.** Protein network of statistically inhibited kinases.

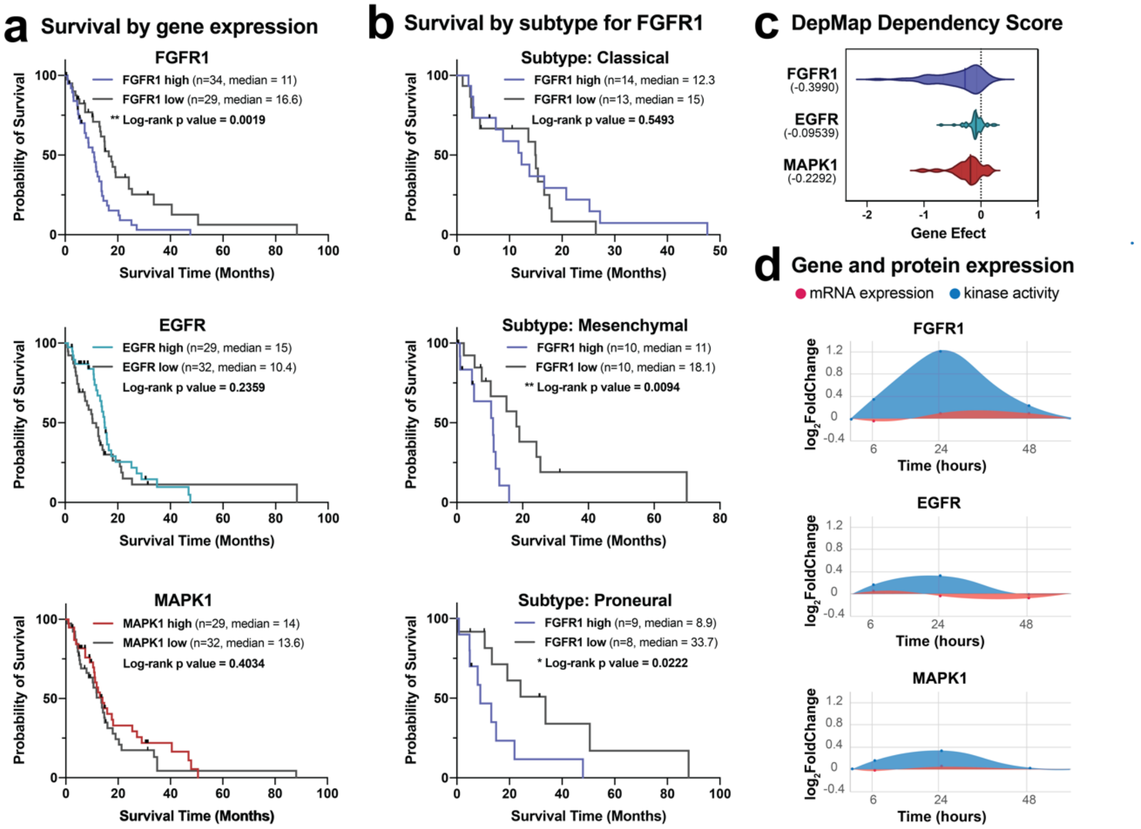


**Supplementary Figure 2. FGFR1 is identified as a top kinase target in GBM. a. Kaplan Meier analysis shows high FGFR1 expression is linked to poor overall survival for GBM patients.** TCGA GBM patients were stratified into high (upper quartile) and low (lower quartile) groups based on RNA-seq gene expression for FGFR1, EGFR, or MAPK1 mRNA levels. Survival was plotted on a Kaplan-Meier curve with a log-rank (Mantel-Cox) test for significance. **b. FGFR1 gene expression is linked to poor overall survival in mesenchymal and proneural GBM subtypes.** TCGA GBM patients were stratified into high and low expression and survival was plotted for classical patients (top), mesenchymal patients (middle), and proneural patients (bottom). **c. Dependency scores show that GBM tumors are highly dependent on FGFR1 for survival.** CRISPR DepMap gene effect scores (version 23Q2) for FGFR1, EGFR, and MAPK1 were calculated for each GBM cell line. A more negative gene effect suggests a higher dependency on the gene for cell survival. **d. Kinase mRNA and protein expression reveals kinase activation in the absence of upregulation.** Kinase activity from MIB/MS screen and mRNA levels from RNA-sequencing was plotted for FGFR1, EGFR, and MAPK1 relative to time-matched DMSO controls.

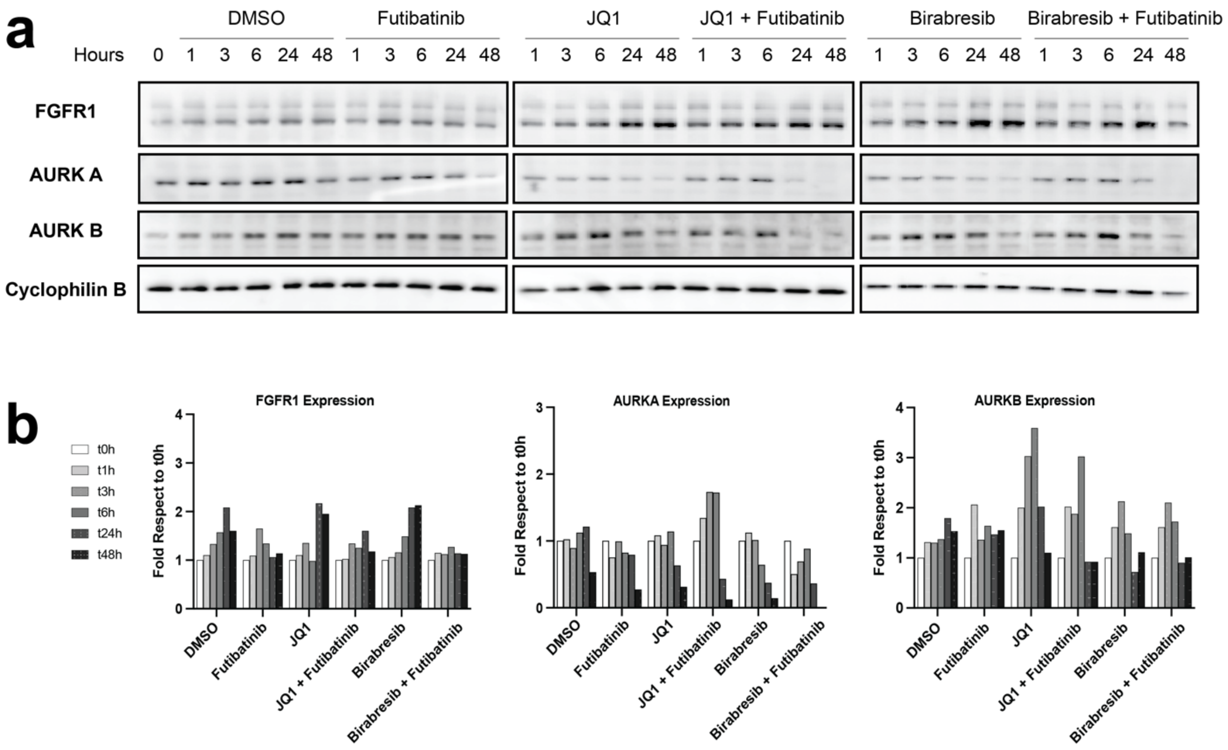


**Supplementary Figure 3. Expression of FGFR1 is increased in GBM22 cells treated with BET inhibitors. Duplicate of Figure 2. a. FGFR1 and Aurora kinases expression.** GBM22 cells treated with DMSO as a control or futibatinib, JQ1, JQ1+futibatinib, birabresib, birabresib+futibatinib, for 0, 1, 3, 6, 24, or 48 hours. Cell lysates were immunoblotted for FGFR1, Aurora Kinase A (AURKA) and Aurora Kinase B (AURKB). Equal loading was verified by immunoblotting for cyclophilin B from the respective gel. For cyclophilin B, the blot shown is from the same blot as FGFR1. The whole experiment was done in triplicate and one representative image is shown. Additional replicates can be found in Supplementary Figures 3-4. **b. Quantification of protein expression from western blots in (a).** Quantification values for the target proteins were normalized to volumes of cyclophilin B loading control and plotted as fold difference from the time zero-hour point from the respective gel.

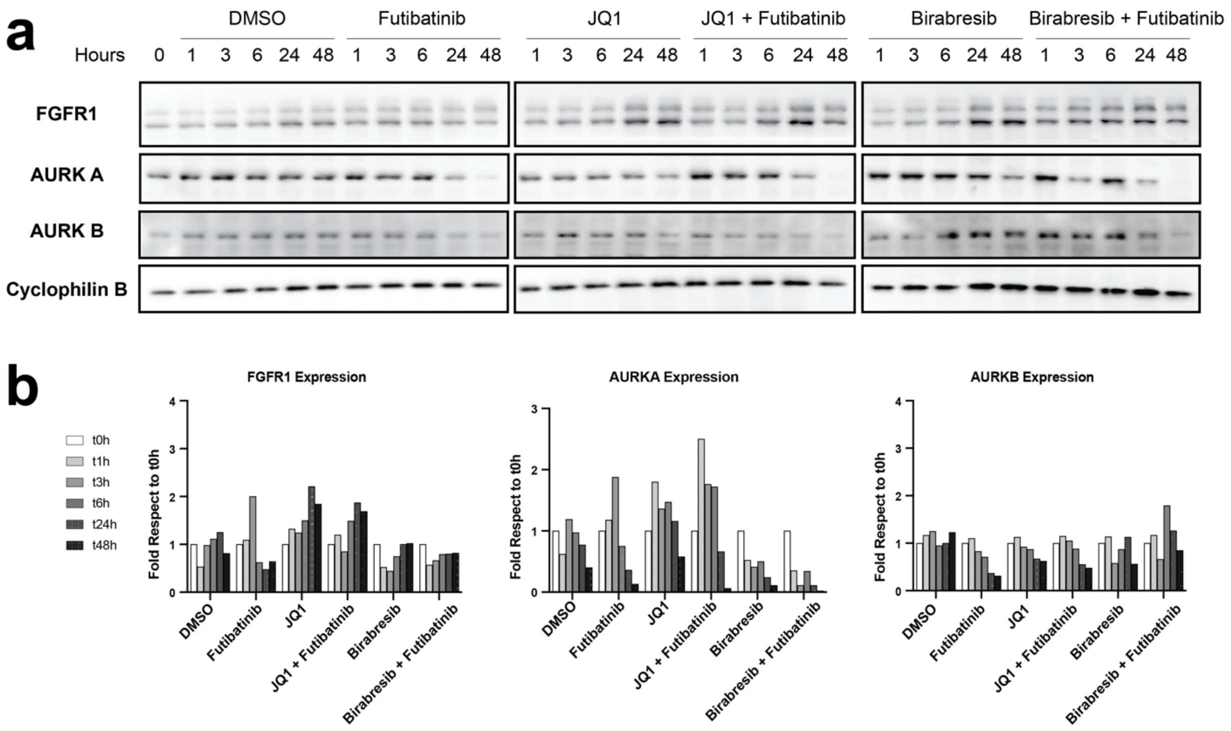


**Supplementary Figure 4. Expression of FGFR1 is increased in GBM22 cells treated with BET inhibitors. Triplicate of Figure 2. a. FGFR1 and Aurora kinases expression.** GBM22 cells treated with DMSO as a control or futibatinib, JQ1, JQ1+futibatinib, birabresib, birabresib+futibatinib, for 0, 1, 3, 6, 24, or 48 hours. Cell lysates were immunoblotted for FGFR1, Aurora Kinase A (AURKA) and Aurora Kinase B (AURKB). Equal loading was verified by immunoblotting for cyclophilin B from the respective gel. For cyclophilin B, the blot shown is from the same blot as FGFR1. The whole experiment was done in triplicate and one representative image is shown. Additional replicates can be found in Supplementary Figures 3-4. **b. Quantification of protein expression from western blots in (a).** Quantification values for the target proteins were normalized to volumes of cyclophilin B loading control and plotted as fold difference from the time zero-hour point from the respective gel.

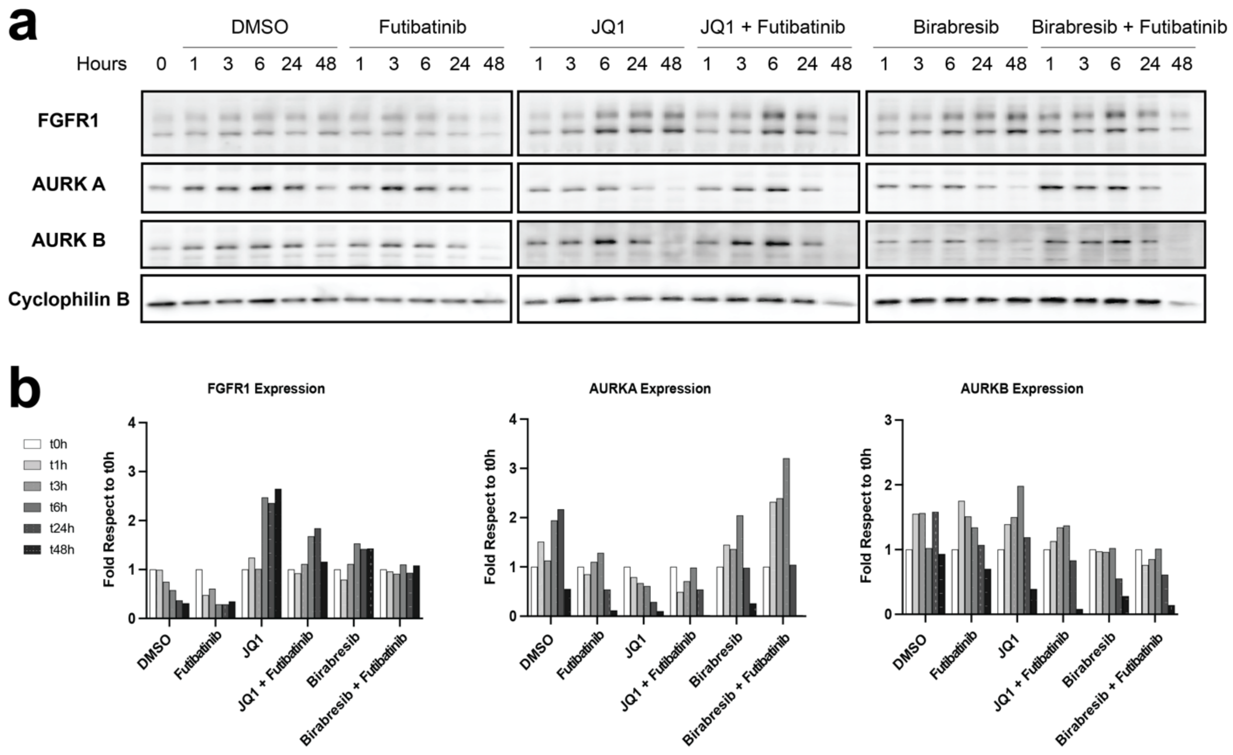


**Supplementary Figure 5. Expression of FGFR1 is increased in GBM6 cells treated with BET inhibition. a.** FGFR1 and Aurora Kinases expression. GBM6 cells treated with DMSO as a control or TAS120, JQ1, JQ1 + TAS120, MK8628, or MK8628 + TAS120, for 0, 1, 3, 6, 24, 48 hours. Cell lysates were immunoblotted for FGFR1, Aurora Kinase A (AURKA), and Aurora Kinase B (AURKB). Equal loading was verified by immunoblotting for Cyclophilin B from the respective gel. For Cyclophilin B, the blot shown is from the same blot as FGFR1. **b**. **Quantification of proteins expression from western blots in (a).** Quantification values for the target proteins were normalized to values of Cyclophilin B loading control and plotted as fold difference from the time zero-hour point (t0h) from the respective gel.

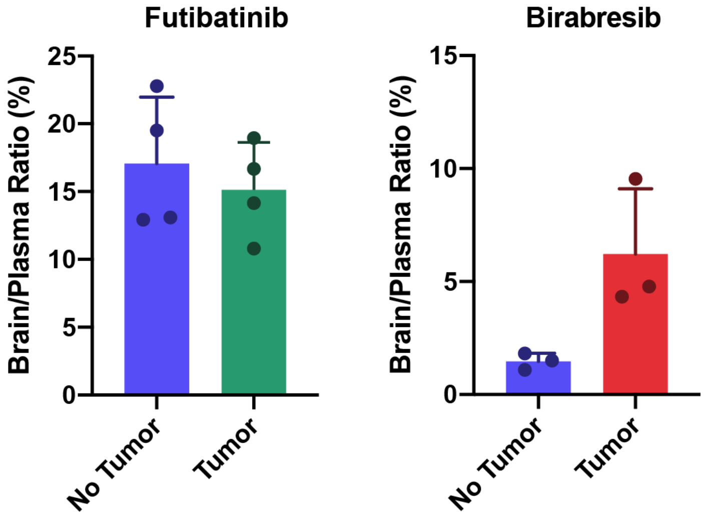


**Supplementary Figure 6. Brain to plasma ratio of drug found in each brain hemisphere using an orthotopic model of GBM.** Six mice were implanted with orthotopic intracranial GBM cells into the right cortex and tumors were established over the course of three weeks. Mice were treated with futibatinib or birabresib by intraperitoneal injection and brains were isolated two hours later. The concentration of drug in each hemisphere was measured using mass spectrometry and the ratio of brain to plasma was plotted for each hemisphere.

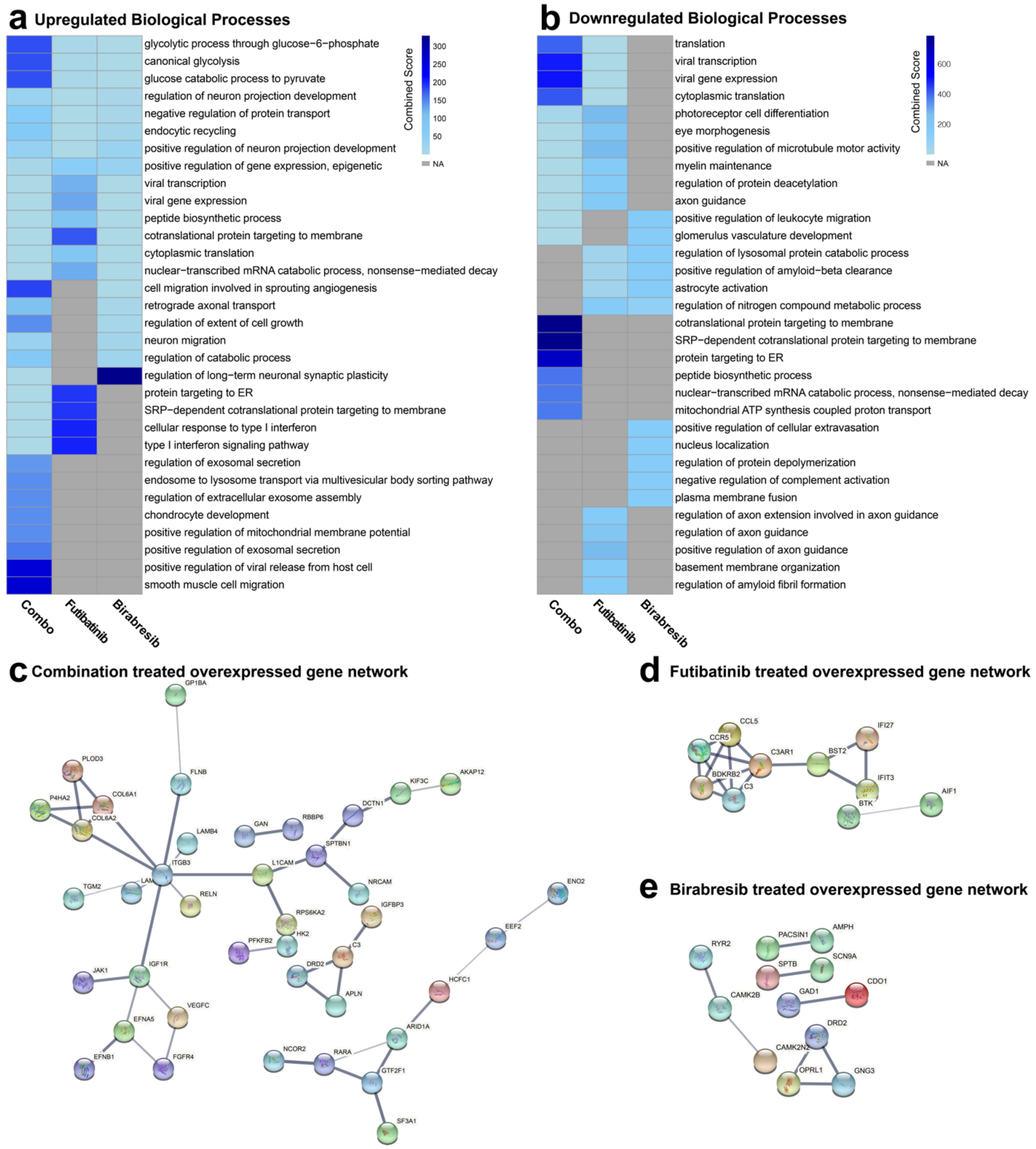


**Supplementary Figure 7. Gene ontology and networks analysis reveals pathways of resistance to combined FGFR and BET inhibition *in vivo*. a-b. Gene ontology of (a) upregulated or (b) downregulated differentially expressed genes reveals enriched biological processes.** Mice with orthotopic GBM39 tumors were treated with futibatinib and/or birabresib and tumors were isolated for bulk RNA sequencing. Genes determined to be differentially expressed by the NOISeqBIO R-package were uploaded to EnrichR to determine human upregulated biological processes. Combined score was calculated as c = log(p) * z, where c = the combined score, p = Fisher exact test p-value, and z = z-score for deviation from expected rank. The top ten upregulated or downregulated processes as determined by combined score for each condition were plotted on a heatmap. NA = not applicable, process was not enriched for this sample. **c-e. STRING protein networks reveal pathways of resistance to combined FGFR and BET inhibition *in vivo*** Mice with orthotopic GBM39 tumors were treated with futibatinib and/or birabresib, then tumors were isolated for RNA sequencing. The top 100 differentially expressed and upregulated genes were input into STRING v11 (https://string-db.or) for pathway analysis and interaction networks.

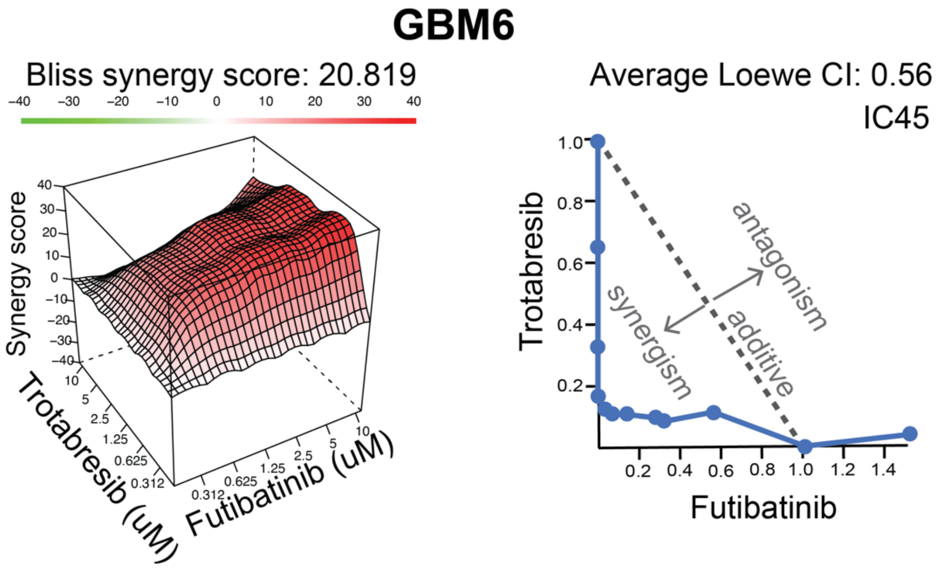


**Supplementary Figure 8. The BET inhibitor trotabresib synergizes with futibatinib *in vitro* in GBM.** GBM6 cells were treated with a dose response matrix of the FGFR inhibitor futibatinib in combination with the BET inhibitor trotabresib. Cell death was measured as the amount of ATP present using CellTiter-Glo®. Bliss independence model synergy plot (left) and Loewe additivity isobologram (right) of the combination indices for each cell line at the indicated inhibition level.
